# Seed-competent tau monomer initiates pathology in PS19 tauopathy mice

**DOI:** 10.1101/2022.01.03.474806

**Authors:** Hilda Mirbaha, Dailu Chen, Vishruth Mullapudi, Sandi Jo Terpack, Charles L. White, Lukasz A. Joachimiak, Marc I. Diamond

## Abstract

Tau aggregation into ordered assemblies causes myriad neurodegenerative tauopathies. We previously reported that tau monomer exists in either inert (M_i_) or seed-competent (M_s_) conformational ensembles, and that M_s_ encodes strains, which are biologically active, self-propagating assemblies. We have previously isolated M_s_ from tauopathy brains, but it is unknown if disease begins with M_s_ formation followed by fibril assembly, or if M_s_ derives from fibrils and is an epiphenomenon. Consequently, we studied a tauopathy mouse model (PS19) that expresses full-length human (1N4R) tau containing a disease-associated mutation (P301S). Using tau repeat domain biosensor cells, we detected insoluble tau seeding activity at 2 months. We found insoluble tau protein assemblies by immunoblot at 3 months. We next immunopurified monomer from mice aged 1-6 weeks using size exclusion chromatography. We detected soluble seeding activity at 4 weeks, before insoluble material or larger assemblies, with assemblies ranging from n=1-3 tau units. By 5 and 6 weeks, large soluble assemblies had formed. This indicated the first detectable pathological forms of tau were M_s_. We next tested for post-translational modifications of tau monomer from 1-6 weeks. We detected no phosphorylation unique to M_s_ in PS19 or Alzheimer’s disease brain. We conclude that tauopathy begins with formation of M_s_ monomer, whose activity is phosphorylation-independent. M_s_ self-assembles to form oligomers before it forms insoluble fibrils. The conversion of tau monomer from M_i_ to M_s_ thus constitutes the first detectable step in the initiation of tauopathy in this mouse model, with obvious implications for origins of disease in humans.

## Introduction

Tauopathies such as Alzheimer’s disease (AD), chronic traumatic encephalopathy, frontotemporal dementias, and related disorders are each characterized by ordered assemblies of the microtubule-associated tau protein (MAPT) (1). We have proposed that transcellular propagation mediated by tau prions underlies their relentless progression (2-7), as have others(8,9). According to this model, aggregates that form in one cell escape to gain entry to connected or adjacent cells, where they act as templates for their own replication. Tau normally binds microtubules(10,11) and does not form detectable aggregates. The initial cause of tau assembly formation in tauopathies is unknown, and it could be that high concentrations of monomer coalesce to form “seeds,” i.e. forms that serve as templates for ordered assembly growth. Indeed, some have proposed that tau undergoes liquid-liquid phase separation(12-15), which could theoretically initiate assembly formation through molecular crowding. By contrast, our prior work has suggested that a seed-competent tau monomer (M_s_) exists in a conformational ensemble that is distinct from inert tau monomer (M_i_) (16,17). Another study is consistent with this idea(18), although this concept is not widely accepted. We have recently developed methods to create M_s_ from M_i_ using recombinant protein (16,19,20), and we have also isolated it from human tauopathies(16,17). Intriguingly, M_s_ adopts multiple seed-competent forms that serve as templates to create defined tau prion “strains” (17), which are tau species that faithfully replicate their structure *in vivo*, and produce defined patterns of neuropathology(5,6,21). We recently proposed that a critical conformational change from M_i_ to M_s_ represents the first step in the aggregation process(16), that M_s_ encodes strains (17), and we have reported methods to produce M_s_ *in vitro*(20). However, it is unknown whether M_s_ precedes or follows the formation of large assemblies *in vivo*—a critical question for the origin of tauopathies.

We previously developed methods to purify full-length tau monomer from brain tissue using a combination of immunopurification and size exclusion chromatography(16). This readily discriminates tau monomer from larger assemblies. We have now used this approach to determine when M_s_ first appears in a transgenic model of tauopathy, and what is its relationship to higher-order assemblies. We also compare tau monomer from AD vs. control patients, and test whether phosphorylation correlates with M_s_ seeding activity.

## RESULTS

### Insoluble tau and seeding activity vs. neurofibrillary pathology

PS19 mice express full-length (1N4R) tau containing a P301S mis-sense mutation that causes dominantly inherited tauopathy in humans(22). They have been extensively characterized by our lab and others, and develop tau neurofibrillary pathology at approximately 6 months of age. We repeated this analysis, staining sections at 1-6 months with AT8(23), an anti-phospho-tau antibody (pS202/pT205), using sagittally sectioned hemibrains (Fig. 1A), and saving the other half for biochemistry (below). In lateral sections neurofibrillary tangles were very rare at 3 months, slightly increased at 4 months, and easily detectable at 6 months. We consistently observed neurofibrillary tangles only at 6 months (Fig.1A), consistent with our prior work(24). Using the contralateral hemi-brain from each animal, we analyzed brain homogenates for soluble tau by western blot (Fig. 1B), and observed no change between 1-6 months. Detergent (sarkosyl) extraction of tau is commonly used to detect fibrillar aggregates, so we compared sarkosyl-insoluble fractions at different ages. We observed insoluble tau by western blot at 3 months (Fig. 1C), several months before the appearance of obvious cellular tau pathology.

**Figure 1.**
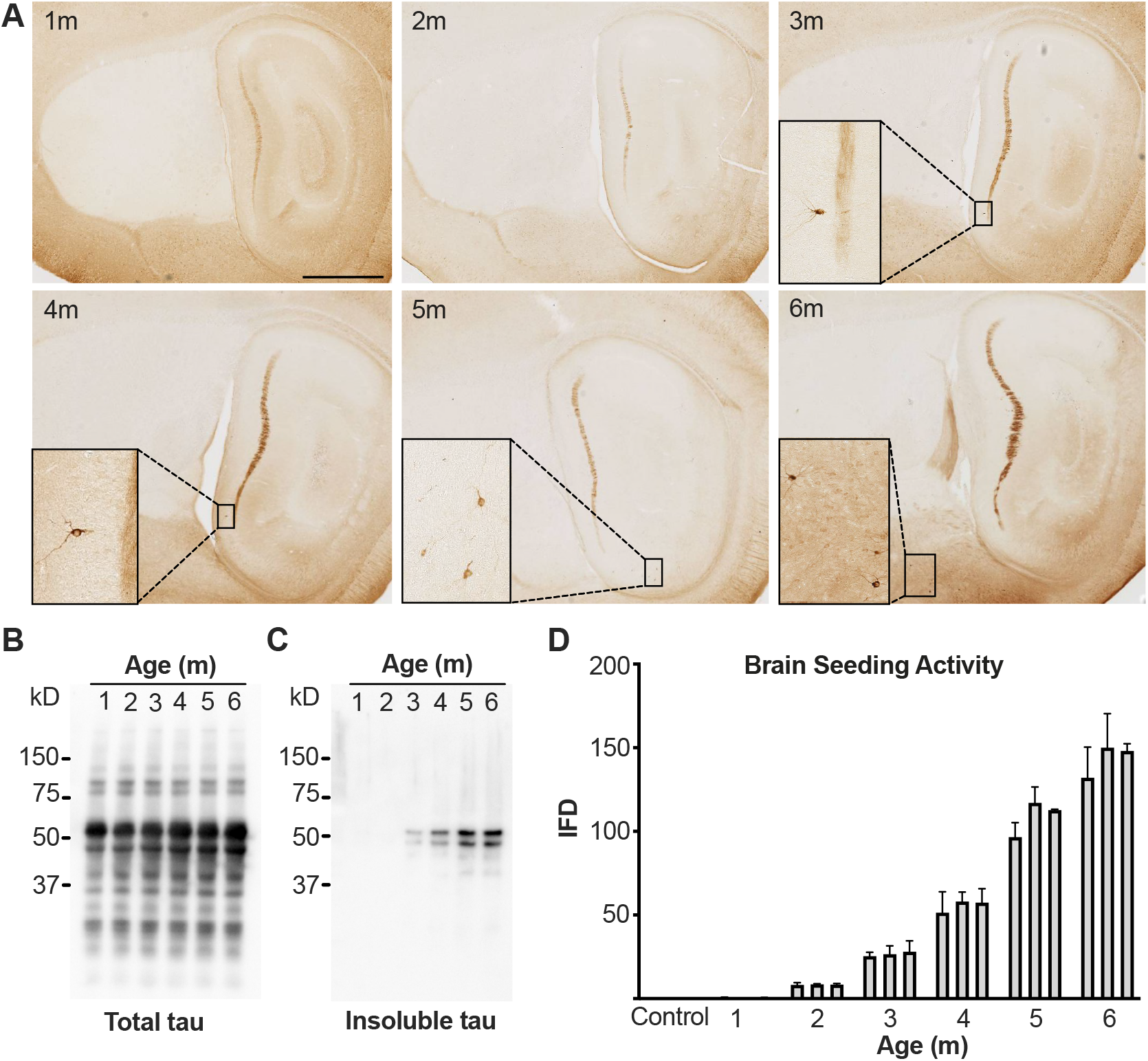
Neurofibrillary pathology and seeding activity of insoluble tau in aging PS19 mice. (**A**) Sagittally sectioned hemibrain of PS19 mice age1-6 months immunostained by AT8 antibody which highlights rare neurofibrillary tangles in 3 months in lateral sections of hippocampus and increases by age (representative images shown; scale bar = 2 mm) In medial sections of hippocampus, neurofibrillary tangles were only observed in 6 months (not shown). (**B**) Western blot of PS19 contralateral hemibrain homogenate of aging mice shows no difference in total tau expression level at all ages from 1 to 6 months. (**C**) SDS-Page analysis of detergent-insoluble tau extracted from PS19 mice brain (3 for each age mixed and loaded on the gel) shows insoluble tau is detectable at 3 months and increases with age. (**D**) Seeding activity (reported as integrated FRET density) of detergent-insoluble tau appears in mice at 2 months and increases with age. Each bar represents one mouse.

To determine when detergent-insoluble seeding activity appears, we tested each of the fractions in a well-established cellular biosensor assay based on expression of tau repeat domain containing a single disease-associated mutation (P301S) fused to cyan and yellow fluorescent proteins(24). We observed no seeding at 1 month, and detected minimal activity at 2 months that grew steadily thereafter (Fig. 1D).

These experiments established that in PS19 mice overall human tau levels are constant between 1-6 months; insoluble seeding activity occurs at 2 months; insoluble tau is apparent at 3 months by western blot, and abundant neurofibrillary pathology does not appear until 5-6 months.

### Soluble tau assemblies appear at 4-6 weeks

Having established the onset of large, insoluble tau assemblies, we extended our study to younger mice ranging from 1-6 weeks (n=3 per age) to characterize precisely the small changes in assembly state from monomer to dimer, trimer, ∼10mer, and ∼20mer. We have previously determined that M_s_ produced *in vitro* exists in dynamic equilibrium with dimers, and potentially trimers(20). We created combined the 3 brains at each time point and extracted sarkosyl soluble and insoluble fractions that we analyzed by western blot. We detected no change in soluble or insoluble tau in this time frame (Fig. 2A). We then used HJ8.5 antibody, which binds the N-terminus of human tau(25), to immunoprecipitate the brain homogenates. We resolved the supernatant from a 21,000 x g centrifugation using size exclusion chromatography with a Superdex 200 column, according to our established methods(16,26). We have previously determined that there is no detectable cross-contamination between fractions that contain multimers and those that contain tau monomer(16). We performed western blots for purified tau using polyclonal anti-tau antibody (Agilent). We observed no larger tau assemblies until 4 weeks of age, and these increased in abundance between 4 and 5 weeks (Fig. 2A). We also used 2µm filter dot blot to characterize the fractions, which confirmed the findings regarding soluble vs. insoluble tau, and the prevalence of small assemblies (Fig. 2B).

**Figure 2.**
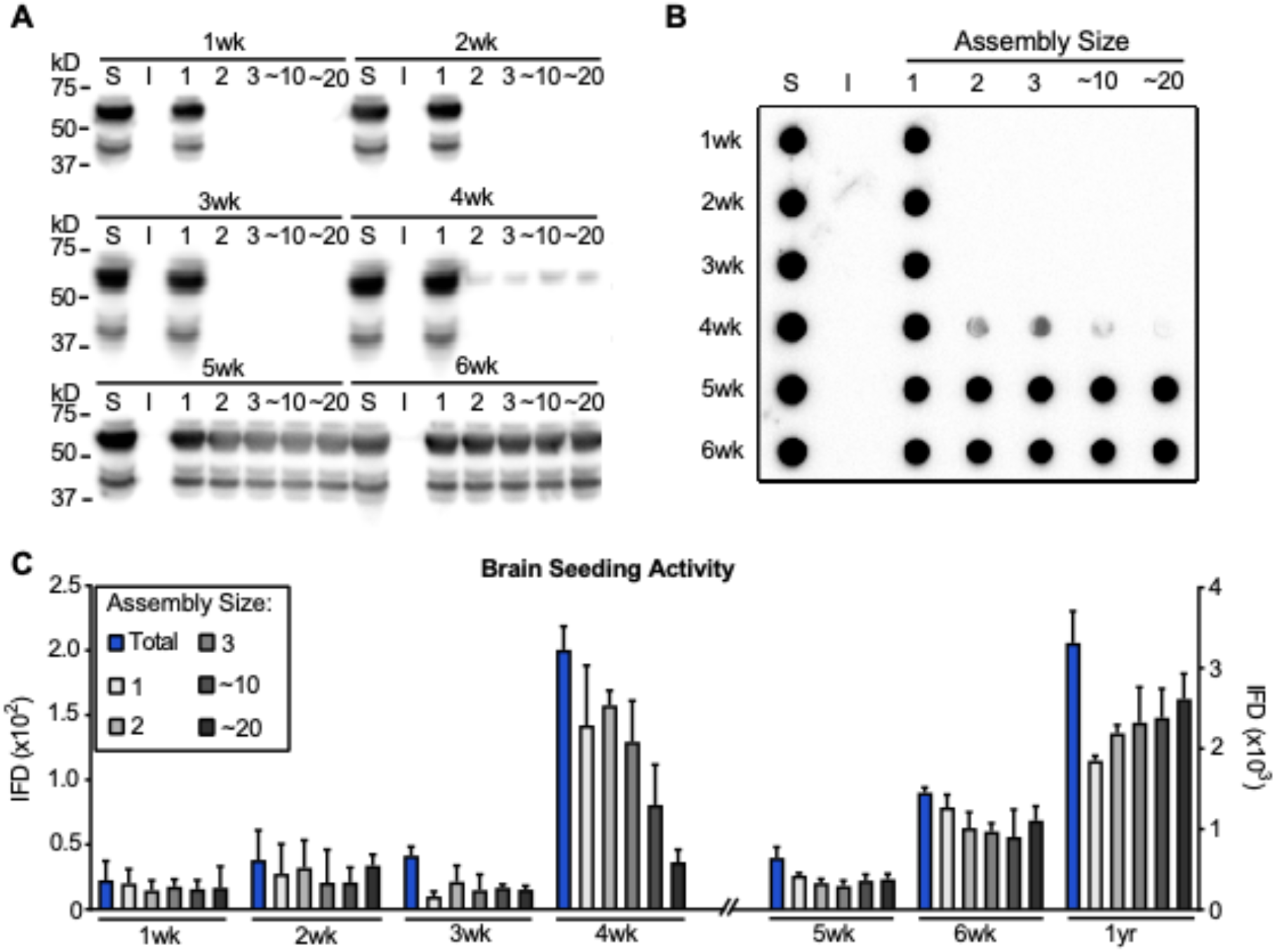
Seeding activity of soluble tau in PS19 mice brain at 1 to 6 weeks. To determine the assembly state of tau before formation of insoluble tau, we purified tau from supernatant after 21,000 x g centrifugation of PS19 brain homogenate from mice age 1-6 weeks with an anti-tau monoclonal antibody (HJ8.5). Immunoprecipitated tau was resolved by size exclusion chromatography. (**A**) SDS-PAGE was used to analyze S=total soluble tau; I=sarkosyl-insoluble tau; or assemblies of various sizes ranging from n=1 to ∼20mer. Tau monomer alone was present until 3 weeks, while assemblies appeared at 4 weeks with increasing abundance by 5 and 6 weeks. (**B**) Similar analyses were carried out using dot-bot. (**C**) The 21,000 x g supernatant (T), and tau assemblies were seeded to RD-CFP/YFP biosensors. Seeding activity was first detected at 4 weeks, with a rapid increase by 5 and 6 weeks. IFD = Integrated FRET Density. Assembly size: 1 = monomer, 2 = dimer, 3 = trimer, ∼10 = ∼10mer, ∼20 = ∼20mer. Error bars=S.D. IFD: integrated FRET density. Note different IFD scales for weeks 1-4 and 5-6, and 1 year. Image of full western blot is in Supplemental Figure 2.

Next we tested the seeding activity of these fractions using biosensor cells. We found no seeding activity through 3 weeks. We detected low seeding activity at 4 weeks, which increased dramatically at 5-6 weeks. At 4 weeks, monomer and smaller size oligomers (n=1-3) accounted for most of the seeding, whereas by 5 weeks and beyond the seeding distributed across a range of assembly sizes (Fig. 2C). These results established that the very first seed-competent forms occur as tau monomer and very small assemblies, whereas larger, detergent-insoluble forms of protein appear several weeks later.

### M_i_ and M_s_ have similar post-translational modifications

It is unknown what initiates the conversion of M_i_ to M_s_, or what might maintain M_s_ in a seed-competent state. Tau phosphorylation was first noted in the setting of tau pathology (27-29) and has been studied for decades. Hence we determined the phosphorylation pattern of tau monomer isolated across mice aged 1-6 weeks, and in tau monomer purified from AD and control patients, and in sarkosyl-insoluble tau from AD patients. After immunopurification, monomer was isolated by size exclusion chromatography, denatured, proteolyzed with trypsin and analyzed by mass spectrometry to determine post-translational modifications. We observed a relatively conserved pattern of tau phosphorylation from week 1 to week 6, and some deviation in the insoluble material from 1 year old mice (Fig. 3A, Supplemental Fig. 3-1A). Phosphorylation patterns observed in tau monomer at 1-6 weeks persisted at 1 year. We observed no phosphorylation changes that explained the rapid increase in M_s_ seeding activity at 3-6 weeks.

**Figure 3.**
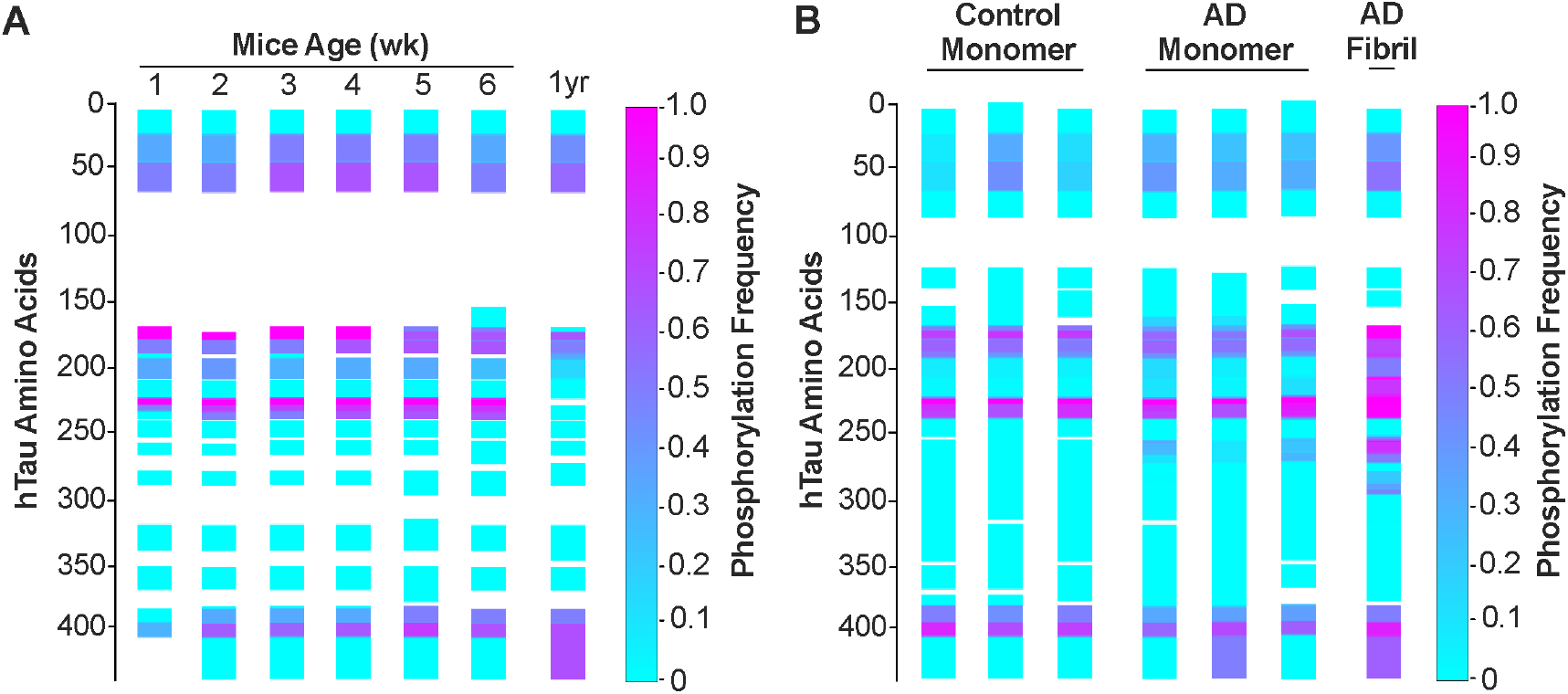
Phosphorylation patterns on tau monomer isolated from aging PS19 mice, AD, and age-matched controls. (**A**) Tau monomer isolated from PS19 mouse brains age 1-6 weeks (n=3 animals per week) was analyzed by mass spectrometry to determine phosphorylation patterns. Intensities of modified and unmodified peptides were compared to calculate and abundance of the modification. Phosphorylation frequency is coded from 0% (cyan) to 100% (magenta). (**B**) Similar analysis was performed on tau monomer samples isolated from 3 age-matched controls and 3 AD patients, and an AD fibril prep, with human peptides mapped against 2N4R tau sequence. Gap in PTM seen in mice reflects 1N4R tau.

We performed similar analyses on tau monomer isolated from human AD and control brain, and also from tau fibrils extracted from AD brain. We observed phosphorylation patterns in control and AD monomer samples similar to mice (Fig. 3B, Supplemental Fig. 3-1B). We observed higher phosphorylation levels in both number of modification sites and their frequencies in the insoluble AD tau fraction (Figure 3B, Supplemental Figure 3-1B, Supplemental Figure 3-2). As for the mouse samples, in human samples we observed no phospho-residues that differentiated AD from control monomer, except S262, which was modified in 4-20% of peptides across the three AD samples but not controls (Supplemental Fig. 3-2). S262 was modified in AD tau fibrils at ∼60% frequency (Supplemental Fig. 3-2). Human tau monomer was highly phosphorylated in all samples at T231, with control at 65-71%, and AD at 82-94%, and ∼100% of AD fibrils positive (Supplemental Fig. 3-2). Overall, we observed no patterns that differentiated AD vs. control monomer.

We also measured tau ubiquitination and acetylation. We detected no modifications consistent with ubiquitination in tau monomer at any age of PS19 mice. We detected low levels of ubiquitination in tau monomer from human control and 2 of the AD brains (Supplemental Fig. 4). K254 was ubiquitinated in all conditions at <1% frequency. Monomer from one AD patient exhibited 3 peptides ubiquitinated at 100% frequency (Supplemental Fig. 3-3). However, ubiquitination frequencies of those residues accountable were mostly <1% except for K257, with 71% ubiquitination. AD tau fibrils exhibited more ubiquitination sites and higher frequencies than monomer. (Supplemental Fig. 3-3). We observed acetylation on K44 in tau monomer at 2-4 weeks in PS19 mice, but all at a very low frequency of 0.4-0.7%. Although peptide 156-170 at 6 weeks showed 100% acetylation at K163, this peptide was not detected in samples of other ages, and therefore no clear conclusion can be derived for this position. In human tau monomer, we observed acetylation on K44 in all 3 control monomer and 2/3 AD at a low frequency of 0.3 - 5%. AD fibrils had acetylation on K375 at frequency of 69% (Supplemental Fig. 3-4). Taken together, we found no correlation of ubiquitination or acetylation with seeding activity in tau monomer of PS19 mice, as ubiquitination was not detected at any age studied, while acetylation occurred at a very low level in both seed-positive and -negative conditions. In human samples, we observed several sites of acetylation and ubiquitination within the repeat domain region of AD fibrils. However the occurrence of acetylation and ubiquitination on tau monomers was low, either from control or AD samples (with one exception of ubiquitination in AD3 monomer). Overall, we did not detect changes in acetylation or ubiquitination of M_s_ that explained its seeding activity.

## DISCUSSION

Tauopathies are defined by fibrillar assemblies of tau protein. Our prior studies have defined two conformational states of tau monomer, inert (M_i_) and seed-competent (M_s_). M_s_ self-assembles to form larger aggregates(16), and also encodes strains in biosensor cell systems(17). A critical unanswered question is whether M_s_ is derived from pre-existing tau fibrils, or whether it forms first as an initial step in pathogenesis, and then leads to larger assemblies. Notably, we have determined that M_s_ typically exists in dynamic equilibrium *in vitro* with dimers and potentially trimers(20). We have now determined that M_s_ appears in the brain of a mouse model (in conjunction with dimers and trimers) within a defined time window, before larger assemblies have formed, and in the absence of detergent-insoluble deposits. We conclude that the first events in neurodegeneration, prior to the development of neurofibrillary tangles, comprise a shift in tau conformation from M_i_ to M_s_. At least in the PS19 mouse model, where we can conduct longitudinal studies, we propose that disease onset is defined at the first moment of conformational change from M_i_ to M_s_.

### No phosphorylation pattern distinguishes M_s_ and M_i_

Post-translational modifications on insoluble tau, especially phosphorylation, acetylation, and ubiquitination, are well-described in humans(31), to the point where modification, especially phosphorylation at S202/T205 is used routinely to classify the extent of disease in humans(32) and mice(33) with the AT8 monoclonal antibody(23). This has led to the logical hypothesis that PTMs play a role in pathogenesis. Our findings do not support a primary role for tau PTM in disease initiation, at least in PS19 mice. When we resolved tau monomer over one week intervals in the PS19 mice, we observed no modifications that correlated with conversion of M_i_ to M_s_, despite an increase in seeding activity of >50-fold in the 4-6 week time window. Further, in MS analyses of tau from human brains, AD monomer (which had seeding activity) more closely resembled that of control brain than insoluble tau from AD, which had many PTMs. MS reports on all tau modifications together, and cannot determine whether a tiny seed-competent fraction of monomer has a specific PTM that is responsible for its activity. To partially address this question, we have tested whether tau monomer derived from PS19 mice (Supplemental Figure 4A,B) or two different AD cases (Supplemental Figure 4C,D) retains its seeding activity after protein phosphatase PP2A treatment. We could not confirm whether every phospho-residue was removed, but we did not observe any loss of seeding activity following phosphatase treatment.

In summary, we cannot exclude that monomer seeding activity at 4 weeks results from a low abundance of distinctly modified peptide not detected by mass spectrometry. Additionally, we have not accounted for any potential variance in the kinase/phosphatase activities during the developmental stages of mice. Nevertheless, the fact that monomer seeding vs. week 4 increased 20-fold at week 5, and 100-fold at week 6, with essentially no change in overall phosphorylation patterns, indicates that within the limits of detection (and in contrast to insoluble tau) we could not correlate tau M_s_ seeding with a distinct phosphorylation pattern.

### M_s_ precedes tau assembly

In our prior work we have observed that M_s_ can be created by sonication of fibrils(16,26), or by conversion of M_i_ following heparin exposure(16,20). Work by the Nussbaum-Krammer lab indicates that chaperone treatment of fibrils *in vitro* liberates seed-competent monomer and possibly tau oligomers(18). Hence in humans and mice it has remained an open question as to which actually comes first: large assemblies vs. M_s_, and whether M_s_ might derive from pre-existing fibrillar species. This work answers this question as well as possible with current technology. We find that in a very short period of time, over a 7 day window between week 3 and 4 of life in a PS19 mouse, M_i_ begins to convert from a state in which it is exclusively monomer to a seed-competent form (with associated dimers and trimers) but very few larger species. Yet by 5-6 weeks, a full range of assembly sizes is detected. This is about 2-6 weeks before the appearance of detergent-insoluble (large) tau assemblies, which we detected by seeding activity at 2 months, and by western blot at 3 months. The simplest interpretation of these results is that seed-competent tau monomer, M_s_, is formed before subsequent oligomers and fibrillar species, and does not derive from them. While not directly addressed here, this observation calls into question the idea that seeds form only after local high concentrations of tau have occurred. Instead, we favor a model in which specific events somehow facilitate conversion of tau from one conformational state (M_i_) to another (M_s_), prior to formation of larger, more stable assemblies.

### Conversion of M_i_ to M_s_ marks initiation of tau pathology *in vivo*

The goal of any mechanism-based therapy for tauopathy will be to intervene before disease onset. Hence a major challenge has been to define the point in human disease when this occurs. We still do not know the answer to this question in humans, but our demarcation of a 7 day period between weeks 3-4 in the life of a PS19 mouse, during which we can detect the first changes in tau conformation, indicate that M_s_ formation precedes all subsequent neuropathology. Hence we propose that the biochemical origin of disease occurs at this moment, and that understanding mechanisms that underlie this process will reveal the basis of neurodegeneration caused by subsequent large tau assembly formation.

## MATERIALS AND METHODS

### Isolation of mouse brain

Animals were anesthetized with isoflurane and perfused with cold PBS. Brains were hemi-dissected. The right hemisphere was frozen in liquid nitrogen and stored at -80°C for subsequent biochemical assays while the left hemispheres were drop-fixed in Phosphate-Buffered 4% Paraformaldehyde (FD NeuroTechnologies, Colombia, MD, USA) overnight at 4 °C. Left hemispheres were then placed in 10% sucrose in PBS for 24 hr at 4 °C, followed by 24 hr in 20% sucrose in PBS at 4°C, and finally stored in 30% sucrose in PBS at 4°C until sectioning.

### Immunohistochemistry of mouse tissue

A sliding-base freezing microtome (Thermo-Fisher Scientific, Waltham, MA, USA) was used to collect 40 μm free-floating sagittal sections from the fixed mouse brains. Sections were stored in cryoprotectant at 4 °C until IHC performed. Slices were first blocked for one hour with 5% BSA in TBS with 0.25% Triton X-100 (blocking buffer). Brain slices were incubated with biotinylated AT8 antibody (1:500, Thermo Scientific) overnight in blocking buffer at 4°C. Slices were subsequently incubated with the VECTASTAIN Elite ABC Kit (Vector Labs, Burlingame, CA, USA) in TBS prepared according to the manufacturer’s protocol for 30 minutes, followed by DAB development using the DAB Peroxidase Substrate Kit (Vector Labs). Slices were imaged using the Olympus Nanozoomer 2.0-HT (Hamamatsu, Bridgewater, NJ, USA) in the Univ of Texas Southwestern Medical Center Whole Brain Microscopy Core Facility, RRID:SCR_017949).

### Tau extraction from mouse/human brain and characterization by SEC

Right hemispheres of 3 PS19 mice brains of identical ages ranging from 1 to 6 weeks and 1 year old were gently homogenized in 1:10 ratio w/v of TBS buffer containing protease inhibitor cocktails (Roche) using a dounce homogenizer. Samples were centrifuged at 21,000 x g for 15 min at 4°C to remove cellular debris. The supernatant loaded onto a Superdex 200 Increase 10/300 GL column (GE Healthcare) and fractions including the monomer and oligomers were partitioned into aliquots, snap frozen and stored at -80°C for seeding assay and western blotting.

For the PTM studies, left hemispheres of 3 mouse brains of the identical ages were dounce homogenized, centrifuged at 21,000 x g to remove cellular debris. The supernatant was collected and incubated with 400µg of monoclonal anti-tau antibody (HJ8.5) and 2ml of Pierce™ ProteinA/G magnetic beads (ThermoFisher) overnight. The beads were washed 3 times and eluted with a low pH buffer and immediately neutralized. The eluted sample were run on a Superdex 200 (10/300) column (GE) to fractionate different tau species including monomer, dimer, trimer, 10-mer and 20-mer. Monomer fraction was concentrated by 5KMWC filter (Pierce), snap frozen and used for mass spectrometry studies of PTMs.

To extract insoluble Tau fibril from mouse brain it was dounce homogenized in PHF buffer (10 mM Tris-HCl (pH 7.4), 0.8 M NaCl, 1mM EDTA, protease and phosphatase inhibitor) at a 1:10 ratio (w/v). The homogenate was then centrifuged at 21,000 x g and 1% sarkosyl was added to the supernatant followed by end-to-end rotation at room temperature for 1 hr. It was then centrifuged at 186,000 x g for 1 hr and the pellet was washed with PBS followed by second round of ultracentrifugation at 186,000 x g. The supernatant was discarded and the pellet was resuspended in 1 mL PBS to be analyzed by seeding assay and western blot.

Tau extraction from human brains was adopted from (16) and is similar to the procedure for mouse brains. 1g of frontal lobe sections from AD patients at late Braak stage (VI) and age-matched controls lacking evident tau pathology were gently homogenized at 4°C in 10 mL of TBS buffer containing protease inhibitor cocktails (Roche) using a dounce homogenizer. Samples were centrifuged at 21,000 x g for 15 min at 4°C to remove cellular debris. Supernatant was partitioned into aliquots, snap frozen and stored at -80°C. Immunopurification was performed with HJ8.5 anti-tau antibody(25) at a ratio of 1:50 (1 µg mAb per 50 µg of total protein), incubating overnight at 4°C while rotating. Beads were washed with TBS buffer before overnight incubation at 4°C. The complexes were centrifuged at 1000 x g for 3 min and the supernatant was discarded.

Beads were washed with Pierce™ Gentle Ag/Ab Binding Buffer, pH 8.0 (ThermoFisher) three times. Tau was eluted from the beads using 180 µL low pH elution buffer (ThermoFisher), incubated at room temperature for 10 min, followed by neutralization with 18 µL Tris-base pH 8.5. The residual beads were removed by magnet, and the supernatant was loaded onto a Superdex 200 Increase 10/300 GL column (GE Healthcare). SEC fractions were flash frozen and stored at -80°C.

### Western and dot blot

For western blot and dot blot analyses, immunoprecipitation was eliminated to maximize the quantity of tau protein and enhance tau visibility on the blots. SEC fractions were boiled for 5 min with SDS-PAGE sample buffer, and loaded into a NuPAGE 4–12% Bis-Tris Gel in a chamber filled with NuPAGE™ MOPS SDS running buffer (ThermoFisher) and run at 100V for ∼110 min. Samples were then transferred to a PVDF membrane using a semi-dry transfer apparatus (Bio-Rad). After being blocked in 5% milk (Bio-Rad), the membrane was incubated with primary anti-tau polyclonal antibody (Dako, Agilent) at 1:2000 on a shaker overnight at 4°C. It was then washed with TBST 3 times and was incubated with anti-rabbit secondary antibody for 1 hr at room temperature. The membrane was then washed with TBST on a shaker for 5 min twice, followed by another wash of TBST for 1h, before the final rinse with TBS. The membrane was then developed with ECL prime western blot detection kit (GE Lifescience) for 3 min before being imaged by a digital imager (Syngene).

### Liposome-Mediated Transduction of Tau Seeds

Stable cell lines were plated at a density of 35,000 cells per well in a 96-well plate. Eighteen hours later, at 50% confluency, cells were transduced with proteopathic seeds. Insoluble tau extracted from mice hemibrains, was diluted in PBS to the final ratio of 1:200 of a hemibrain homogenate volume (5uL insoluble tau extract + 5 uL PBS per each well of a 96-well plate). This volume was selected based on titrating insoluble tau extract of the 6 month old mice (data not shown) which demonstrated the highest seeding activity withought causing toxicity on biosensor cells. Transduction complexes were made by combining [8.75 μL Opti-MEM (Gibco) +1.25 μL Lipofectamine 2000 (Invitrogen)] with [Opti-MEM + proteopathic seeds] for a total volume of 20 μL per well. Liposome preparations were incubated at room temperature for 20 min before adding to cells. Cells were incubated with transduction complexes for 48 hr.

### FRET Flow Cytometry

Cells were harvested with 0.05% trypsin and then fixed in 2% paraformaldehyde (Electron Microscopy Services) for 10 min, then resuspended in flow cytometry buffer. The MACSQuant VYB (Miltenyi) was used to perform FRET flow cytometry. To measure CFP and FRET, cells were excited with the 405 nm laser, and fluorescence was captured with a 405/50 nm and 525/50 nm filter, respectively. To measure YFP, cells were excited with a 488 laser and fluorescence was captured with a 525/50 nm filter. The integrated FRET density, defined as the percentage of FRET-positive cells multiplied by the median fluorescence intensity of FRET-positive cells, was used for all analyses(24,34). For each experiment, 20,000 cells per replicate were analyzed and each condition was analyzed in triplicate. Data analysis was performed using FlowJo v10 software (Treestar).

### Sample Preparation for Mass Spectrometry and Analysis of Phosphorylation

The sarkosyl insoluble sample as well as the concentrated soluble monomer fractions were denatured in 8M urea and reduced with 2.5mM TCEP at 37°C, 600RPM for 30min. Sample was then alkylated with 5 mM iodoacetamide for 30 min at RT and protected from light. The sample solutions were diluted to 1 M urea with 50 mM ammonium hydrogen carbonate and trypsin (Promega) was added at an enzyme-to-substrate ratio of 1:50. Proteolysis was carried out at 37 °C overnight followed by acidification with formic acid to 2% (v/v). Samples were then purified by solid-phase extraction using Sep-Pak tC18 cartridges (Waters), dried, and the resulting peptides were reconstituted in 10 uL of 2% (v/v) acetonitrile (ACN) and 0.1% trifluoroacetic acid in water. A portion of the sample was injected onto an Orbitrap Fusion Lumos mass spectrometer (Thermo Electron) coupled to an Ultimate 3000 RSLC-Nano liquid chromatography systems (Dionex). Samples were injected onto a 75 μm i.d., 75-cm long EasySpray column (Thermo), and eluted with a gradient from 0-28% buffer B over 90 min. Buffer A contained 2% (v/v) ACN and 0.1% formic acid in water, and buffer B contained 80% (v/v) ACN, 10% (v/v) trifluoroethanol, and 0.1% formic acid in water. The mass spectrometer operated in positive ion mode. MS scans were acquired at 120,000 resolution in the Orbitrap and up to 10 MS/MS spectra were obtained in the ion trap for each full spectrum acquired using higher-energy collisional dissociation (HCD) for ions with charges 2-7. Dynamic exclusion was set for 25 s after an ion was selected for fragmentation. Raw MS data files were analyzed using Proteome Discoverer v2.2 (Thermo), with peptide identification performed using Sequest HT searching against the human or mouse protein database from UniProt, including the tau isoform of interest. Fragment and precursor tolerances of 10 ppm and 0.6 Da were specified, and three missed cleavages were allowed. Carbamidomethylation of Cys was set as a fixed modification, with oxidation of Met and phosphorylation of Ser, Thr, and Tyr set as a variable modifications. The false-discovery rate (FDR) cutoff was 1% for all peptides. An in-house Python script was created (Github link: https://git.biohpc.swmed.edu/s184069/abundanceparser) to parse out the relative abundance (in percentage) of the phosphorylated, acetylated and ubiquitinated peptides to the total abundance for each peptide sequence detected. An in-house Matlab script was generated to illustrate the relative abundance of PTM peptides with color-coding on a sequence map of modified peptides.

### Recombinant PP2A production

Genes encoding the full-length sequences of Human PP2A A subunit and PP2A C subunit with N-terminal His tag and non-cleavable HA tag were sub-cloned into the pFastBac-Dual vector (Invitrogen). Sequences of PP2A A and C subunits are shown in Table 1. The plasmid containing PP2A A and C subunits was transformed into DH10Bac *E. coli* for transposition into the bacmid. The purified bacmid with Cellfectin II reagent was transfected into Sf9 cells grown in media SF-900 III SFM (Gibco) supplemented with 10% FBS incubated at 27°C with 130 rpm for 7 days. P1 virus was spun down at 1,500 rpm for 5 min and high titer P2 virus stock was produced from the P1 virus (1:200 v/v infection). High Five^™^ insect cells for the expression of PP2A A/C complex were cultured in EX-CELL 405 Serum-Free medium (Millipore-Sigma) at 27°C with 130 rpm. Cells were harvested 2 days after viral infection (1:50 v/v) and resuspended in 50mM Tris-HCl, 100mM NaCl, 2mM MgCl_2_, 5mM βME, and 20mM imidazole, pH8. Lysate was homogenized by GEA Niro Soavi’s PandaPLUS 2000, followed by 15,000 x g spin. The supernatant was incubated with Ni-NTA beads with head-to-head rotation at 4°C for 1hr. The Ni-NTA mix was pelleted at 1000 x g, 4°C for 5min and the pellet resuspended in lysis buffer and loaded onto a gravity column. The bead bed was wash by 40 CV of 50mM Tris-HCl, 100mM NaCl, 2mM MgCl_2_, 5mM βME, and 50mM imidazole, pH8. Although only the PP2A C subunit has His-tag, the high binding affinity of the PP2A A and C subunits allows co-elution of the complex, which was eluted in 50mM Tris-HCl, 100mM NaCl, 2mM MgCl_2_, 5mM βME, and 250mM imidazole, pH8. The Ni elution was buffer exchanged into 20mM Tris-HCl, 50mM NaCl, 2mM MgCl_2_, and 1mM DTT, pH8, by PD-10 column (GE). Exchanged fractions were applied to a HiTrap Q HP (GE) and eluted with a 50 mM–1 M NaCl gradient. Fractions that contain both A and C subunits were collected, aliquoted and flash frozen.

### PP2A dephosphorylation of mouse and human brain isolated tau monomer

Tau monomer from 1-year-old P301S mouse and human AD brain was extracted using the same protocol as described above except for the elimination of immunoprecipitation before SEC and 2-fold higher concentrating for the SEC fraction, which were to enrich tau for the downstream western blot analysis. 50µL of tau monomer from either mouse or human brain was incubated with 50µL of either 10µM PP2A A+C complex or buffer (20mM Tris, 100mM NaCl, 1mM DTT, pH8) for 3hr at 37°C. Immunoprecipitation with 50µL of magnetic Dynabeads™ Protein A (ThermoFisher) was used for each condition to extract tau from the mixture. Beads were first incubated with 5µg of HJ8.5 antibody for 40min followed by a 200µL PBST wash. 100uL of the experiment mixture was diluted 2-fold then incubated with the beads-antibody conjugate for 1.5hr. The beads-antibody-tau conjugate was washed 3 times with 200uL Pierce™ Gentle Ag/Ab Binding Buffer, pH 8.0 (ThermoFisher), after which it was eluted with 45µL Pierce™ IgG Elution Buffer (ThermoFisher) for 5min followed by neutralization with 4.5µL Tris-base pH 8.5. The elution was split for seeding assay and western blots. Seeding assay was conducted following the protocol described above and quantified by FRET flow cytometry. Conditions with or without PP2A were analyzed by SDS-PAGE western blot with rabbit polyclonal anti-tau (Dako, Agilent) to show that the amount of tau after immunoprecipitation was comparable between the two conditions. Zn^2+^ SuperSep™ Phos-tag™ (50μmol/l) 12.5% acrylamide SDS-PAGE (Fujifilm) was used to demonstrate PP2A dephosphorylation efficacy on tau. The phos-tag western blot was used similarly to conventional SDS-PAGE western blot, except for first the use of a different running buffer (25mM Tris, 192mM glycine, 0.1% (w/v) SDS), the use of a special EDTA-free ladder WIDE-VIEW^™^ Prestained Protein Size Marker III (Fujifilm), and an additional step of 3 washes with 10mmol/L EDTA in transfer buffer before the transfer.

## Key resources table

Cell line

Mouse Brain

Human Brain ID

Antibody

Recombinant protein tau sequence

Recombinant protein PP2A A+C sequence

## ACKNOWLEDGEMENTS

We would like to thank Dr. Byung-cheon Jeong and Dr. Xuelian Luo for help with PP2A protein production. Thank Sofia Bali for providing an in-house Matlab script. Research was supported by: Aging Mind Foundation (M.I.D.); Glick Family Foundation (M.I.D.); Cure Alzheimer’s Foundation (M.I.D.); NIH 1RF1AG065407 (M.I.D., L.A.J.); 1R01AG059689 (M.I.D.), 1R56AG061847 (M.I.D.); Brightfocus (L.A.J.); CurePSP (L.A.J.).

## SUPPLEMENTAL FIGURES

**Supplemental Figure 2.**
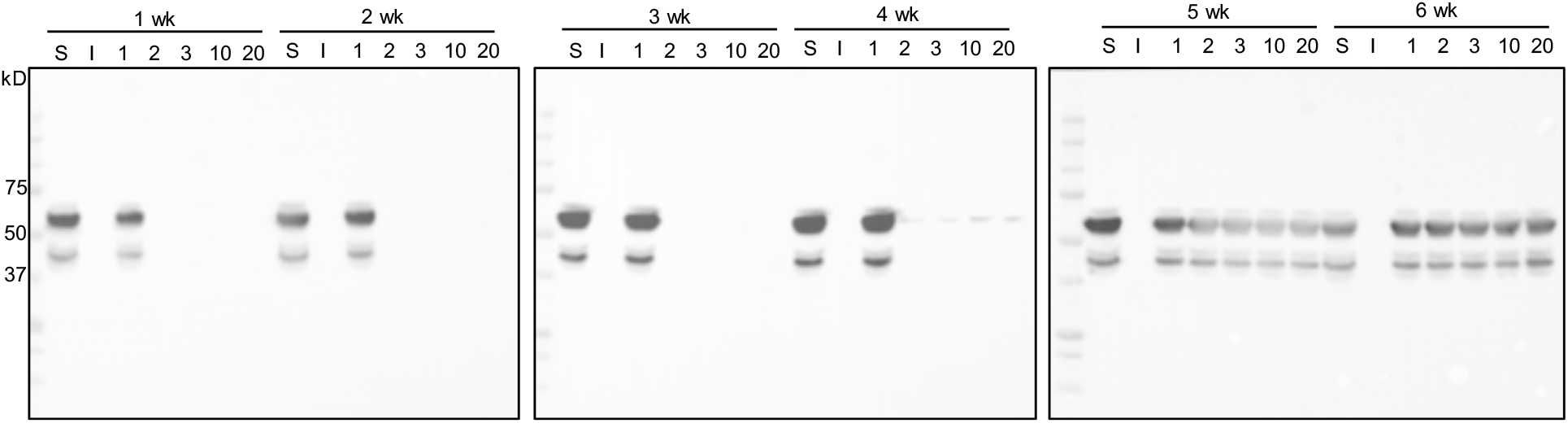
Full western blot image of cropped gel in Figure 2.

**Supplemental Figure 3-1.**
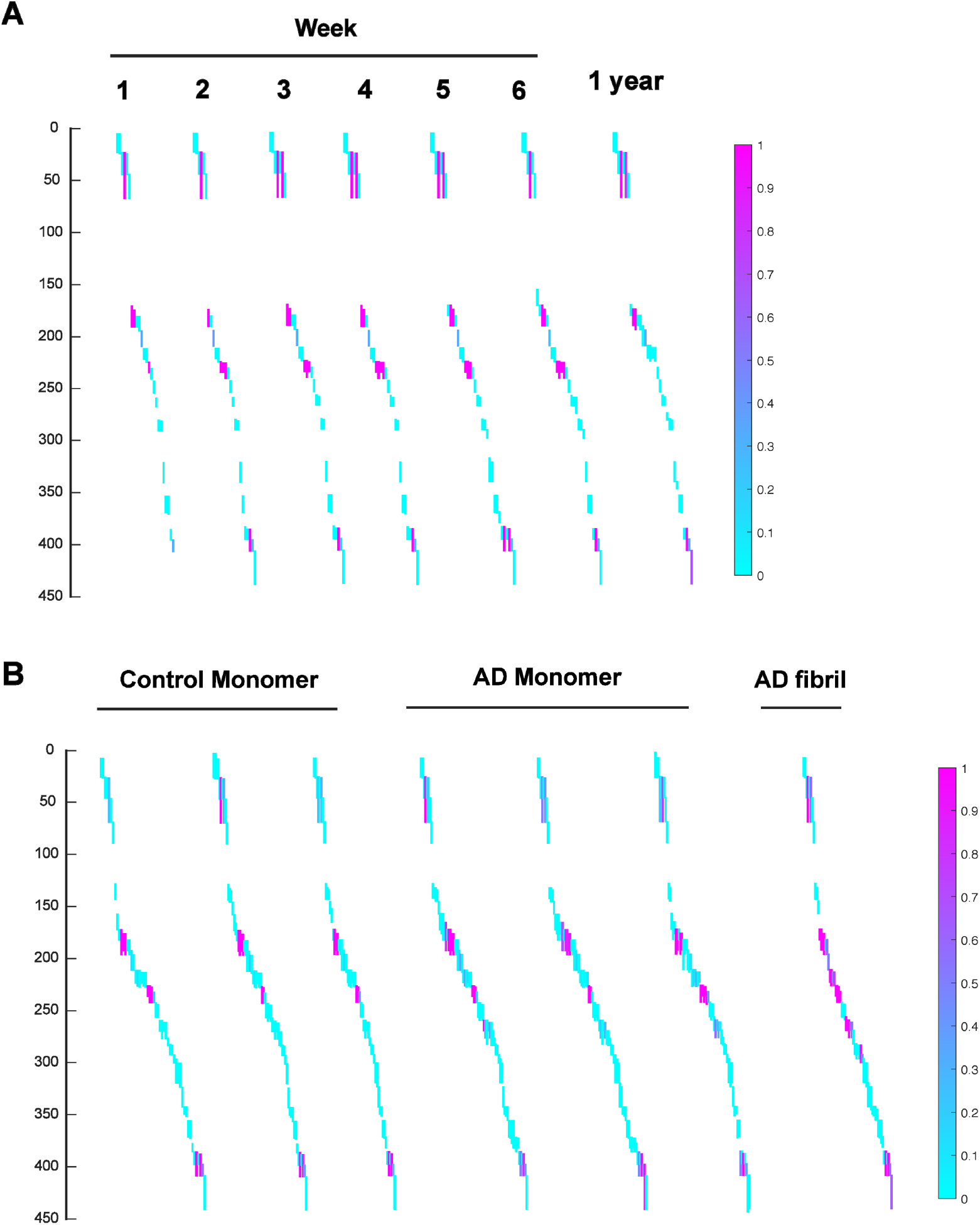
Phosphorylation frequency and peptide coverage for PS19, control monomer, AD tau monomer and AD fibril. (**A**) Peptide coverage map for tau monomer isolated from PS19 mice 1-6 weeks and 1 year. Peptide fragments are colored by phosphorylation frequency from 0 (cyan) to 100% (magenta). (**B**) Peptide coverage map for tau monomer isolated from age-matched controls, AD and AD fibrils. Peptide fragments are colored by phosphorylation frequency from 0 (cyan) to 100% (magenta). Each bar represents a different peptide detected, offset to allow discrimination.

**Supplemental Figure 3-2.**
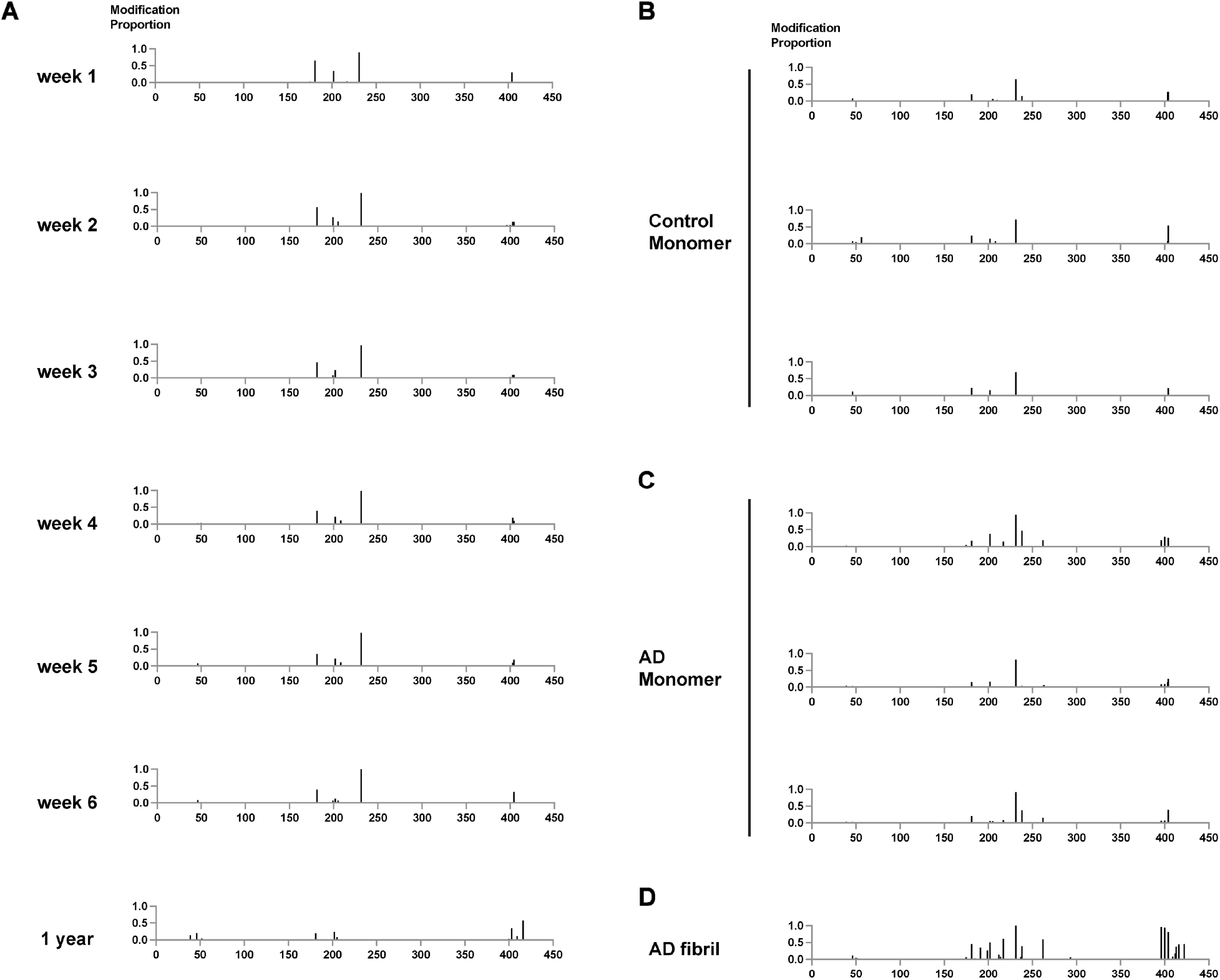
Cumulative phosphorylation frequency by amino acid position. (**A**) Cumulative frequency of phosphorylation for specific sites across different peptide sequences for PS19 tau monomer samples isolated from mice age 1-6 weeks and 1 year. Cumulative frequency of phosphorylation for specific sites across different peptide sequences for tau monomer isolated from age-matched controls (**B**) and AD (**C**) as well as AD fibrils (**D**).

**Supplemental Figure 3-3.**
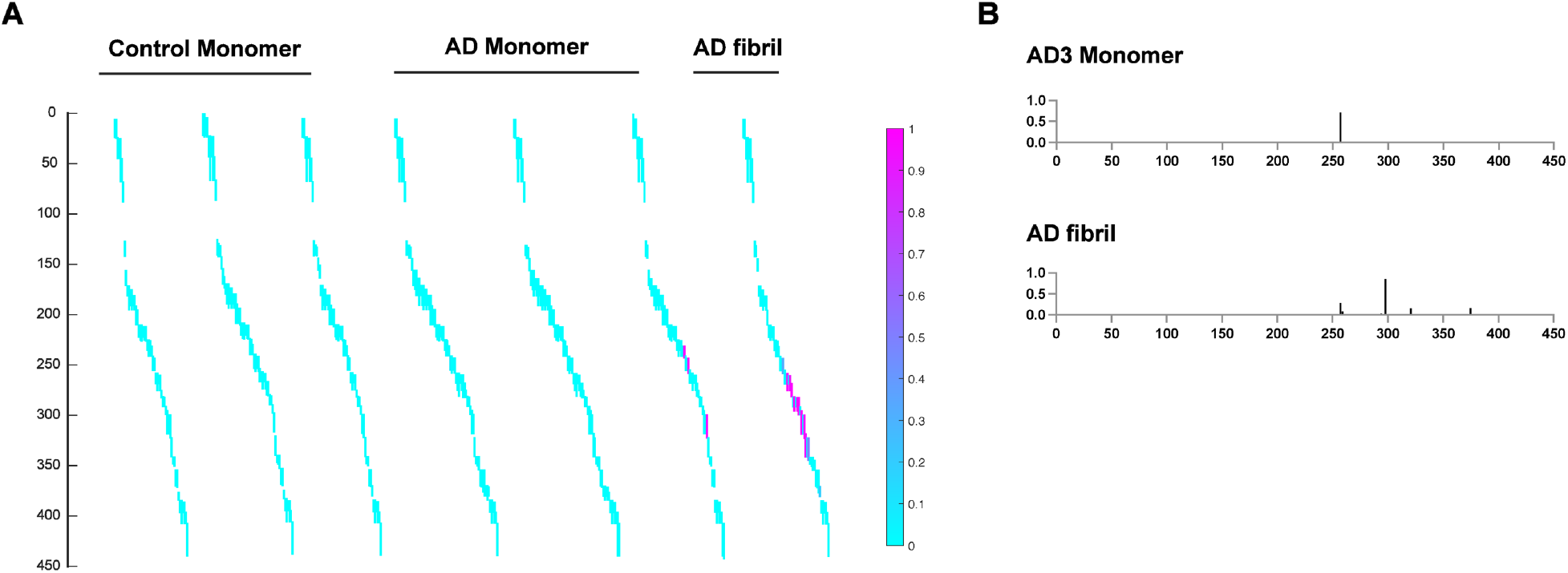
Ubiquitination patterns in AD and control tau samples. (**A**) Peptide coverage map for tau monomer isolated from age-matched controls, AD and AD fibrils. Peptide fragments are colored by acetylation frequency from 0 (cyan) to 100% (magenta). (**B**) Cumulative frequency of ubiquitination for specific sites across different peptide sequences for AD3 tau monomer (the only sample with ubiquitination) and AD fibrils. Each bar represents a different peptide detected, offset to allow discrimination.

**Supplemental Figure 3-4.**
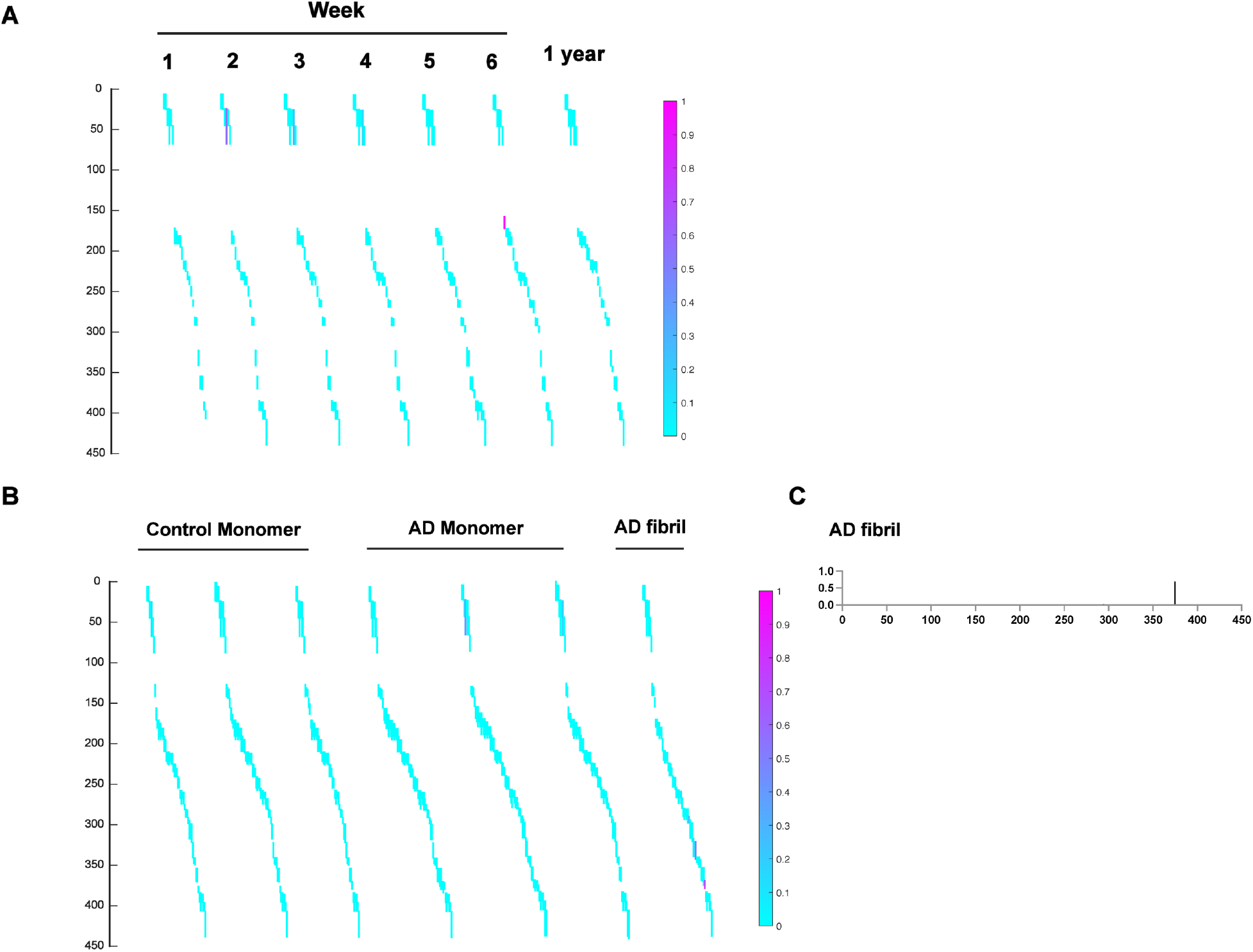
Acetylation patterns for tau monomer isolated from PS19 mouse and human control and AD samples. **(A)** Peptide coverage map for tau monomer isolated from PS19 mice 1-6 weeks and 1 year. Peptide fragments are colored by acetylation frequency from 0 (cyan) to 100% (magenta). (**B)** Peptide coverage map for tau monomer isolated from age-matched controls, AD and AD fibrils. Peptide fragments are colored by acetylation frequency from 0 (cyan) to 100% (magenta). **(C)** Cumulative frequency of acetylation for specific sites across different peptide sequences for AD fibrils. Each bar represents a different peptide detected, offset to allow discrimination.

**Supplemental Figure 4.**
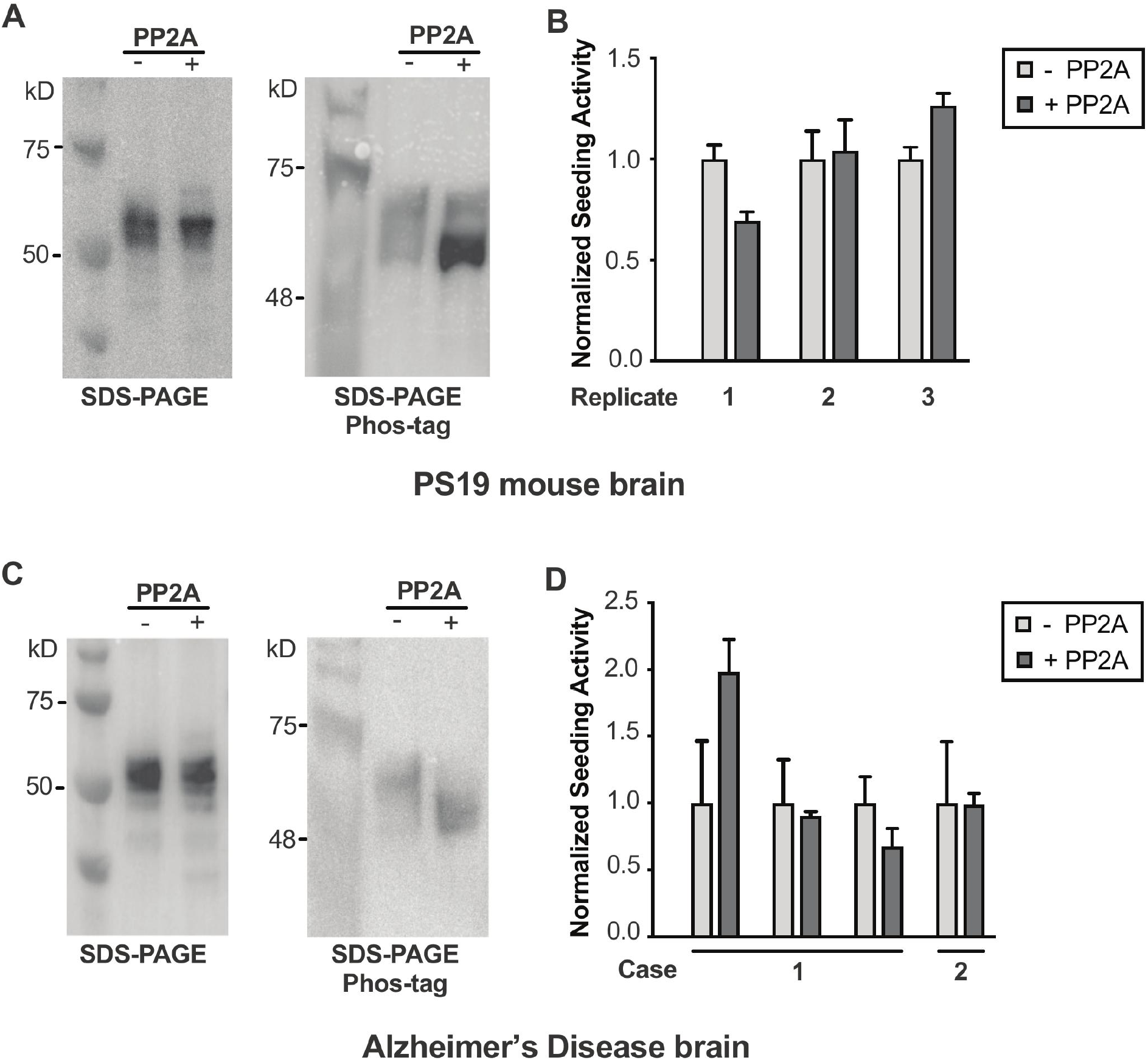
Seeding activity of dephosphorylated tau monomer from AD and mouse. (**A**) Western blot analysis of tau monomer isolated from PS19 mice resolved by SDS-PAGE and Phos-tag^™^ SDS-PAGE probed with anti-tau antibodies reveal patterns consistent with tau phosphorylation at multiple sites. Treatment of samples with phosphatase collapses bands on Phos-tag^™^ gel, indicating dephosphorylation. (**B**) Four independent replicates of tau monomer extracted from 1 year old PS19 mouse brains. Samples were treated with PP2A phosphatase (dark grey) and compared to non-treated comtrol samples (grey) in a tau seeding assay. Error bars=S.D. (**C**) Western blot analysis of tau monomer isolated from AD brains and resolved by SDS-PAGE and Phos-tag^™^ SDS-PAGE probed with anti-tau antibody. Treatment of samples with phosphatase collapses bands on Phos-tag^™^ gel, indicating dephosphorylation. (**D**) Four independent replicates of tau monomer were extracted from 2 different AD brains. Samples were treated with PP2A phosphatase (dark grey) and compared to non-treated samples (grey) in a tau seeding assay. Error bars = S.D.

